# SARC028 samples reveal an interplay between TGF-beta, interferon signaling and low HLA class I expression as contributors to Ewing sarcoma checkpoint blockade resistance

**DOI:** 10.1101/2025.03.26.645492

**Authors:** Jessica D. Daley, Elina Mukherjee, David Ferraro, Shanthi Bhaskar, Anthony Green, Ernest M. Meyer, Hussain Tawbi, Melissa Burgess, Tullia C. Bruno, Anthony R. Cillo, Kelly M. Bailey

## Abstract

**Purpose:** Ewing sarcoma, in contrast to some adult sarcoma subtypes, generally does not respond to single agent immunotherapy targeting PD1. The features of Ewing sarcoma that preclude the effectiveness of immunotherapy remain largely unknown. To address this question, we utilized biopsies from patients with Ewing sarcoma obtained pre- and post-pembrolizumab (anti-PD1) therapy from the phase 2 clinical trial SARC028 to interrogate the Ewing tumor microenvironment and features associated with resistance to checkpoint inhibition.

**Experimental Design:** We utilize multiplexed immunofluorescence, spatial proteomics and spatial transcriptomics to analyze paired pre- and 8 weeks post-treatment biopsy specimens from patients with Ewing sarcoma enrolled on SARC028.

**Results:** Pembrolizumab therapy did not alter the quantity of immune cell infiltration in Ewing tumor biopsies. Analysis of tumor-associated protein markers revealed increased immunoregulatory markers after pembrolizumab. Spatial transcriptomics identified ten cellular neighborhoods (CN) across patients consisting of specific cell subsets. CN10 was consistently observed across patients with poor response. This cellular neighborhood was enriched for a tumor subpopulation with high TGF-β response, low interferon (IFN) response, and low HLA class I expression. IFN response, HLA class I expression, and overall immune infiltration were correlated.

**Conclusions:** Analyses of the tumor microenvironment from Ewing sarcoma biopsies reveals an immunosuppressive triad, the disruption of which should be pursued to improve antitumor immunity. This work highlights the unique insight that can be gained from the analysis of paired patient Ewing sarcoma tumor biopsy samples from clinical trials.

**Statement of translational relevance:** Single agent checkpoint inhibitors have demonstrated limited clinical benefit in bone sarcomas, including Ewing sarcoma (EwS). While *in vitro* and limited patient tumor analyses have suggested mechanisms of immunotherapy resistance in EwS, tumor biopsies following checkpoint inhibitor administration have not been examined. SARC028 was a phase II study investigating blockade of programmed cell death-1 (PD-1) in patients with relapsed/refractory sarcomas, including EwS. Tumor biopsies were collected prior to and 8 weeks after anti-PD1 therapy. Here, we leverage a unique opportunity to understand EwS immunotherapy resistance through analysis of paired tumor samples from SARC028. Using a multi-omics approach, we identify features of EwS that contribute to primary resistance to anti-PD1 including TGF-β, reduced interferon signaling and low HLA class I expression. This study provides an improved understanding of tumor characteristics driving poor response to immunotherapy in relapsed EwS and will guide the development of future trials to modulate this immunosuppressive axis.

## Introduction

Ewing sarcoma (EwS) is an aggressive primary bone tumor that primarily occurs in adolescents and young adults (1). EwS is molecularly characterized by a translocation between members of the FET and ETS families of proteins, most commonly EWSR1::FLI1 or EWSR1::ERG (2). Patients with localized EwS have a 5-year overall survival of more than 80%, based on most recent cooperative group trials (3). However, patients with metastatic or relapsed EwS continue to have very poor outcomes (<30% survival rate) despite clinical trials testing various chemotherapy combinations, surgery, and/or radiation therapy (4). Outcomes for this patients with upfront metastatic or relapsed EwS have not improved in decades, and novel strategies and approaches for treatment of these patients are urgently needed.

Ewing tumor immunobiology, and the effect of immunomodulatory therapies on the Ewing tumor microenvironment (TME), remains relatively unstudied due to the low incidence of EwS and the corresponding paucity of patient tumor samples. Despite these challenges, it has recently been shown that the prevalence of immune subpopulations in the EwS TME may be predictive of survival (5). Our group previously interrogated the tumor immune infiltrates in Ewing tumors at baseline and at the time of disease relapse (6). We demonstrated that Ewing tumor immune infiltration in relapsed tumors is higher compared to tumors at initial diagnosis, and intracellular communication analyses identified CD14+CD16+ macrophages as drivers of immune infiltration (6), suggesting that there may be a role for immunomodulation in the treatment of Ewing sarcoma. Other pre-clinical studies have led to the investigation of immune modulation to treat EwS, such as recent trials investigating the effect of priming NK cells to resist the immunosuppressive cytokine TGF-β on anti-tumor immunity (NCT05634369) (7, 8). Further pre-clinical work is needed to identify and prioritize new immunotherapeutic strategies to disrupt the immunosuppressive TME of relapsed EwS.

Immunomodulatory agents alone or in combination have been widely tested and utilized in clinical trials across many cancer types (9, 10). SARC028 (NCT02301039) was a phase 2 clinical trial of patients 16 years and older investigating the use of single agent pembrolizumab, a PD-1 checkpoint inhibitor, in relapsed/recurrent sarcomas (11), including a cohort of patients with bone sarcomas. Previous quantitative analysis of immune cell populations and PD-1/PD-L1 expression identified that the density of tumor-associated immune cells was predictive of objective response (by RECIST criteria) across sarcoma subtypes that achieved an objective response (12). It has also been demonstrated that in undifferentiated pleomorphic sarcoma (UPS), dedifferentiated liposarcoma (DDLPS), and leiomyosarcoma (LMS), a sarcoma immune classification (SIC) is predictive of response to pembrolizumab therapy (13). However, most sarcoma subtypes included in SARC028 had no objective response to single agent pembrolizumab therapy, including patients with relapsed Ewing sarcoma. Given the varied response to these agents, there has increasingly been a focus on better understanding the mechanisms underlying either response or resistance to immunotherapy (14, 15).

SARC028 included thirteen patients with relapsed/refractory Ewing sarcoma. Importantly, patients on this trial consented to undergo tumor biopsy both pre- and post (8 weeks +/− 1 week)-treatment. On-therapy tumor biopsy specimens such as these are a rare and invaluable tool to improve our understanding of the EwS TME. Here, we performed multiplex immunofluorescence imaging, spatial proteomics, and spatial transcriptomics on EwS biopsy specimens obtained both pre- and post-pembrolizumab therapy on the SARC028 clinical trial to investigate the longitudinal effect of checkpoint inhibition on the EwS TME. We found that treatment with anti-PD-1 led to upregulation of immunoregulatory proteins post-treatment, suggestive of a pharmacodynamic effect and demonstrated that anti-PD-1 therapy is not inert in EwS. We also identified distinct spatially resolved cellular neighborhoods (CN), one of which was consistently found across patients with progressive disease. This immunotherapy resistance-associated cellular neighborhood harbored a tumor subpopulation correlated with reduced immune cell infiltration and that demonstrated low levels of interferon (type I and II) signaling, low levels of human leukocyte antigen class I and enhanced TGFβ signaling. Overall, we have identified an immunosuppressive triad in relapsed Ewing sarcoma that is associated with primary resistance to anti-PD-1 therapy. This finding sheds light on potential combinatorial therapeutic opportunities worthy of future preclinical investigation to modulate this immunosuppressive triad in Ewing sarcoma.

## Material and Methods

### Patient demographics and sample accrual

Institutional review board (IRB) approval was obtained from The University of Pittsburgh STUDY23040147 and a data use agreement was established with the Sarcoma Alliance for Research Through Collaboration (SARC; DUA00004275). The SARC028 clinical trial (NCT02301039) was a phase 2 non-randomized trial to assess the safety and efficacy of pembrolizumab (anti-PD-1 antibody) monotherapy in patients with advanced sarcomas. Tumor biopsies were obtained prior to therapy initiation and after 8 weeks (+/− 1 week) on therapy. Pembrolizumab (200mg IV) was administered every 3 weeks. Thirteen patients diagnosed with EwS were enrolled on trial. Additional optional biopsies were permitted at the time of disease progression; no patients with Ewing sarcoma submitted additional optional biopsy material.

### Generation of tissue microarray

A Tissue Micro Array (TMA) was constructed by the University of Pittsburgh Health Sciences Core Research Facility Pitt Biospecimen Core by expert research histologists utilizing a Beecher MTA-1 Manual Tissue Arrayer, (Estigen, Tartu, Estonia) with a 1mm diameter coring needle. The TMA was designed with 37 cores of de-identified paraffin embedded tissues placed into a new blank paraffin block. This included 21 pre-treatment core specimens from 12 patients and 16 post-treatment core specimens from 9 patients. In general, the biopsy specimens obtained were very small, single cores. To maximize available tissue for analysis on the TMA, more than one core punch was included from patient samples with sufficient material. Imperfections in the TMA such as tears or artifacts were rare. Following construction, the TMA block was incubated overnight at 40°C to ensure adequate adherence of the cores. The TMA was then sectioned utilizing a standard histology microtome, following a 4 µm histology sectioning protocol and subsequently stained.

### Mulitplex immunofluorescence staining of tumor specimens and image acquisition

Multiplexed IHC analysis was performed as described previously (16). Briefly, slide mounted formalin-fixed, paraffin-embedded tumor tissue was deparaffinized. AR6 (Akoya Biosciences, Catalog No. AR600250ML) was utilized for heat induced antigen retrieval in serial cycles to stain tissues for CD68 (1:800, Cell Signaling Technology, Catalog No. 76437, RRID:AB_2799882) and FOXP3 (1:250, Cell Signaling Technology, Catalog No. 12653, RRID: AB_2797979), and AR9 (Akoya Biosciences AR900250ML) was utilized for heat induced antigen retrieval in serial cycles to stain tissues for CD4 (prediluted, Biocare Medical LLC, Catalog No. API3209AA), CD8 (1:200, Biocare Medical LLC, Catalog No. ACI3160A) and CD20 (1:200, Leica Biosystems, Catalog No. NCL-L-CD20-L26, RRID:AB_563521), using Opal detection fluorophores (Akoya Biosciences, Catalog No. NEL811001KT) at the following concentrations: Opal-520 diluted 1:100, Opal-570 diluted 1:150, Opal-540 diluted 1:100, Opal-690 diluted 1:100, Opal-620 diluted 1:100, and Opal-650 diluted 1:150) Cell nuclei were also stained with DAPI (40, 6-diamidino-2-phenylindole). Immunofluorescence images were captured on the Vectra (Perkin Elmer) and analyzed using Phenochart and InForm software.

### Profiling with GeoMx Protein Assay

The TMA slide was prepared following Bruker (formerly, Nanostring) protocols without modification (Manual 10150-04 Manual Slide Preparation). Standard immunohistochemistry protocols were used to visualize morphological markers with SYTO13 for DNA, and anti-NKX2.2, anti-CD68 and anti-CD8 antibodies. Quantitative protein detection utilized antibodies with a UV-photocleavable fluorescent barcode containing a unique molecular identifier (UMI). The TMA was scanned using the GeoMx 4 laser system to generate images at single cell resolution. Regions of Interest (ROI) with a maximum size of 600 μm x 600 μm were generated to include tumors based on NKX2.2 positivity (**Figure S1**). Forty-eight distinct ROIs were created in strategic locations with the goal of capturing as much of the sample material as possible. Though rare, areas of imperfection in the TMA (e.g., tears) were avoided when possible during ROI selection. Individual segments were identified by CD68 and CD8 positivity, as well as non-positive stromal areas, and collected separately. Protein probes were sequentially collected by photocleaving the oligonucleotides from the tissue and collected in separate wells of a 96 well plate. Collected probes were barcoded, processed, and counted on the Bruker/Nanostring nCounter system (Manual 10089-08 nCounter Readout) without modification. Count files were downloaded from the GeoMx and subsequentially indexed to the ROI from the tissue scans for analysis.

### Processing of GeoMx proteomic data and identification of differentially expressed proteins

To normalize the GeoMx data, we established an approach to control for the different ROI sizes and correspondingly different numbers of cells within these regions. To achieve this, we first normalized the GeoMx protein data per region imaged by dividing each analyte expression level per region by the sum of the analytes in that region and then multipling by a scale factor of 1000. We then log-transformed (using the natural log) these normalized counts and used them for downstream analyses. To identify differentially expressed proteins, we performed Wilcoxon rank sum tests and a two-sided alpha of less than 5% was considered significant.

### Provenance and processing of single cell transcriptomic data

Two single-cell RNAseq (scRNAseq) datasets from the TME of human patient EwS tumors were aggregated to construct a reference atlas. The first EwS dataset was enriched for CD45+ immune cells sorted from the tumor microenvironment (6) while the second dataset consisted of all cells from the tumor microenvironment (GSE261693; currently private with a reviewer token available). Together, these datasets provide an in-depth single-cell atlas of the EwS TME. To process these data, we combined the count matrices from all samples into a unified Seurat object. Next, we performed SCTransform-based normalization of the highly variable genes following by reciprocal principal component analysis-guided data integration to control for potential technical confounding variables between individual datasets (17, 18). Next, we performed graph-based clustering on the principal components from the integrated datasets as well as UMAP-based dimensionality reduction from the same principal components for visualization (19). Principle components were heuristically selected based on the cumulative variance explained by each principal component.

### Profiling with CosMx Cancer Transcriptome Atlas

We performed imaging-based spatial transcriptomics analyses of a TMA constructed from EwS biopsy samples derived from the SARC028 clinical trial (11) using the CosMx Human Universal Cell Characterization RNA Panel (1000-plex, Bruker/Nanostring). We also included B2M/CD298, PanCK, CD45, CD3 and DAPI as morphological markers. Following generation of data, cells were segmented, and transcripts were allocated to cells from images and quantified using the manufactureer’s standardized pipeline. We next read the output files into Seurat with the LoadNanoString function. After reading data into Seurat, we next jointly embedded our Ewing sarcoma single-cell RNAseq atlas and the CosMx spatial transcriptomic. This was achieved by performed first filtering the scRNAseq data to only genes that were also present in the CosMx Human Universal Cell Characterization RNA Panel, then performing SCTransform and reciprocal PCA-based integration between the cells derived from spatial transcriptomics and those derived from scRNAseq. After jointly embedding these two types of data, we then perform graph-based clustering of the combined single-cell RNAseq and CosMx datasets. Since we had previously annotated cells in the Ewing scRNAseq atlas, we were then able to perform a label-transfer approach to infer cell types in the CosMx data from the scRNAseq reference dataset, thus efficiently and robustly labeling the cell types in the CosMx dataset. Briefly, cells from the CosMx data that co-clustered with known cell types from the scRNAseq data were considered to be the same cell type, and the inferred label from the scRNAseq cells was transferred to the CosMx cells.

### Cell Neighborhood Analysis of CosMx spatial transcriptomic data

We sought to identify in an unsupervised manner the organizational patterns of cells in space in each biopsy from the SARC028 TMA. To achieve this, we leveraged the annotated cell types and their physical locations within tissues and then created a proximity vector consisting of the proportion of cells presented within a given radius of the cell of interest. Specifically, we choose a diameter of 300 units as the distance with which to enumerate neighboring cell types. We performed these measurements across all cells in the dataset and created a matrix of the vector of cells in the region surrounding each individual cell. Using this matrix, we performed k-means clustering to identify recurrent neighborhoods of cells based on the derived proximity vectors. These inferred spatial cellular neighborhoods were then used for downstream analyses.

### Statistical analysis

For differentially expressed proteins in the GeoMx platform and differentially expressed genes from the CosMx platform, we used Wilcoxon rank sum tests. Nominal two-sided type I error rates of less than 5% were considered significant for the protein data, while a false discovery rate of 5% was considered statistically significant for the CosMx transcriptomic data. Given the small size of this patient tumor cohort (Ewing sarcoma) in the SARC028 clinical trial, this study is not statistically powered to conduct formal comparisons between immune signatures and treatment response.

### Code and data availability

Single-cell RNAseq data is derived from publicly available data deposited in the Gene Expression Omnibus. Code is publicly available for this study (Github).

## Results

### Multiplex immunofluorescence analysis demonstrates that pembrolizumab therapy does not alter density of Ewing immune cell infiltrates

Due to the rarity of Ewing tumor specimens, we sought to maximize the output from the limited tissue obtained from patients with Ewing sarcoma treated on SARC028. A tissue microarray (TMA) of biopsy specimens from 13 patients (**Figure 1A**) was generated. In total, 37 tissue sections were included on the TMA. In total, 8 patients had paired pre- and post-therapy (8 weeks +/− 1 week on therapy) tissues samples available, and of these patients 7 had available disease response data. No cases of pseudo progression were observed in patients with Ewing sarcoma. Utilizing this tissue microarray, we conducted analysis with multiplex immunofluorescence, spatial proteomics, and single-cell imaging-based spatial transcriptomics to determine changes in the quantity of immune cell infiltration, immunoregulatory protein expression, and to define cellular neighborhoods (**Figure 1B**).

**Figure 1.**
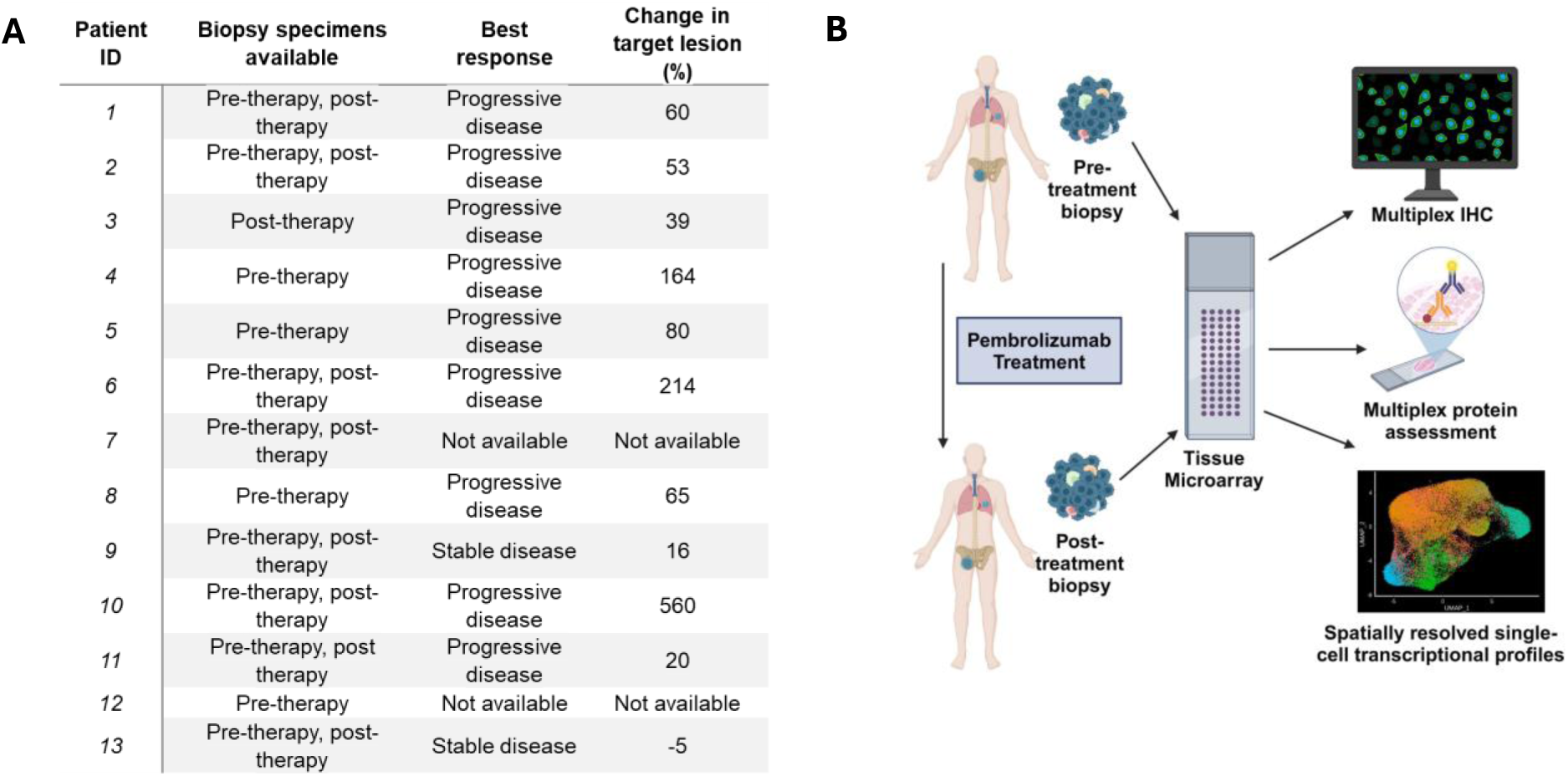
Analysis of Ewing sarcoma biopsy samples pre and post pembrolizumab therapy. **A)** Workflow for generation of tissue microarray (TMA) of Ewing sarcoma biopsy specimens on SARC028 pre and post (8 weeks +/− 1 week) pembrolizumab therapy. TMA was then used to perform multiplex IHC, multiplex protein quantification, and spatially resolved single-cell transcriptional analysis of biopsy specimens. **B)** Patient demographic data of biopsy samples utilized for analysis.

Previous work has demonstrated that the degree of tumor immune infiltration at baseline can be predictive of response to pembrolizumab therapy (15), and tumors with response to pembrolizumab demonstrate increase in immune infiltration following therapy (20). To determine if pembrolizumab altered the quantity of immune cell infiltration in Ewing tumor biopsy specimens, the TMA was analyzed utilizing multiplex immunofluorescence for CD4 and CD8 (T cell markers), CD68 (macrophage marker), CD20 (B cell marker), FoxP3 (regulatory T cell marker), CD45 (pan-leukocyte marker), and DAPI (nuclear stain) (**Figure 2A**). Analyses demonstrate that pembrolizumab therapy did not significantly alter the quantity of total immune cell infiltration in Ewing tumors when comparing all pre-therapy samples to all post-therapy samples (**Figure 2B**). However, on an individual patient level, we did observe both increases and decreases in the quantity of immune cell infiltration following pembrolizumab therapy (**Figure 2C**). The effect of pembrolizumab on the density of immune cell subtypes including CD4+ T cells, CD8+ T cells, regulatory T cells (Foxp3+), B cells (CD20+), as well as macrophage populations (CD68+) was also determined. Pembrolizumab therapy did not significantly alter the total quantity of these immune cell sub-populations in Ewing tumor biopsy specimens following pembrolizumab therapy (**Figure S2**).

**Figure 2.**
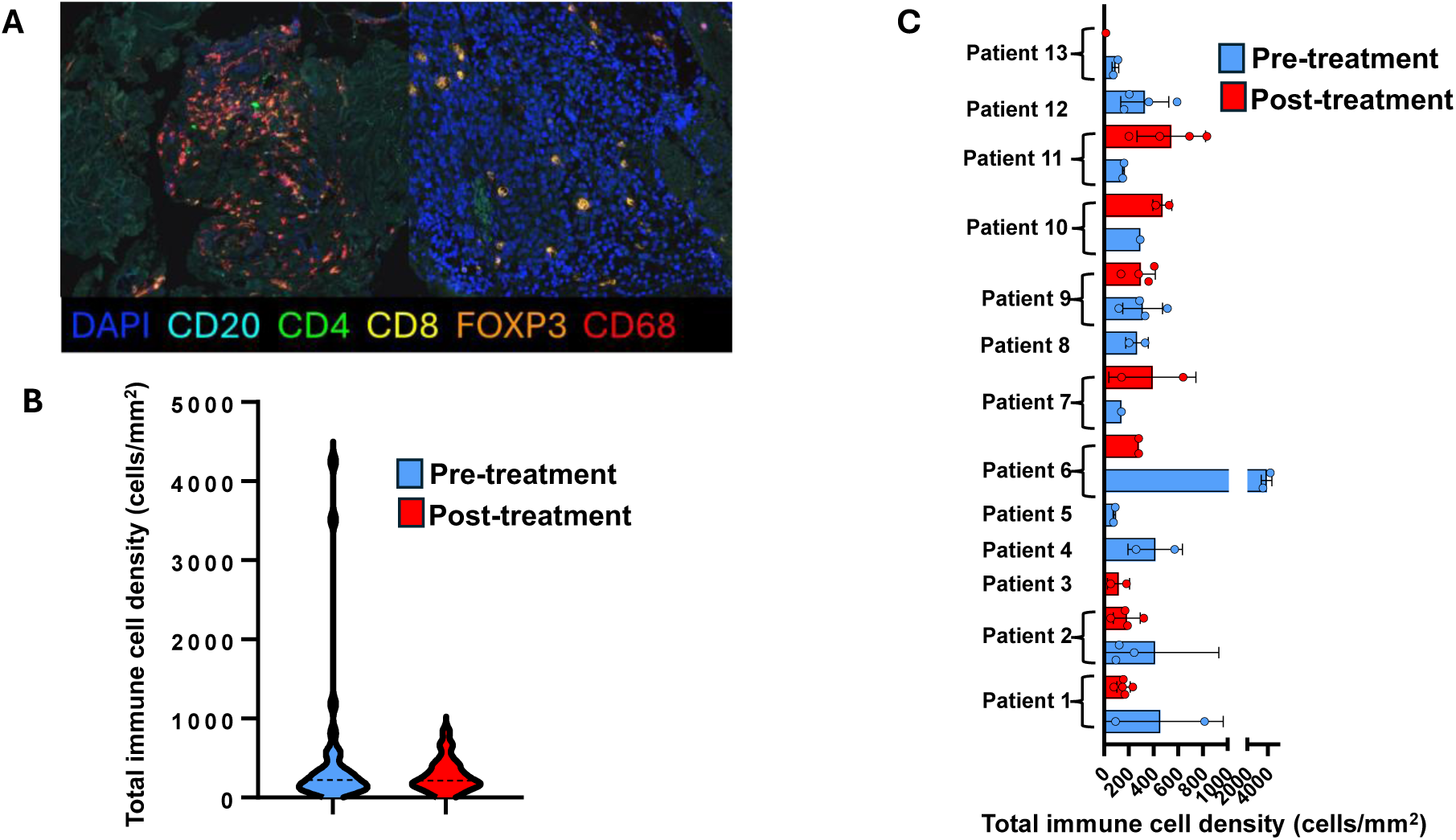
Pembrolizumab therapy does not alter Ewing sarcoma total immune cell density. **A)** Multiplex immunofluorescence for CD4, CD8, CD68, CD20, FoxP3, and DAPI on all tumor biopsy samples. Representative image is shown. **B)** Quantification of total immune cell infiltrate following multiplexed IHC analysis in tumor biopsy samples from 13 patients is shown as an average pre-treatment and post treatment. **C)** Total immune cell infiltrate in each individual sample is shown.

We next sought to investigate if an increase or decrease in the quantity of total immune cell infiltration following pembrolizumab therapy correlated with stable disease or degree of tumor progression in individual patients. In the 7 patients with available paired pre- and post-therapy biopsy samples and tumor response data, analyses did not demonstrate a correlation between change in total immune cell density (all CD45+ cell types) and percent change in tumor size (**Figure S3**). We did observe a positive correlation between the percent change in tumor size and the fold change in both CD68+ cell density and CD4+ cell density (**Figure S3**). However, given the limited number of patient samples, no definitive results regarding tumor response and change in Ewing tumor immune cell infiltration can be concluded. These findings do suggest that a potential reason for the inefficacy of pembrolizumab as a single agent for this patient population was the inability of pembrolizumab alone to significantly increase the quantity of immune cell infiltration in Ewing tumors (versus overall low immune cell infiltration resulting in low efficacy of pembrolizumab).

### Spatial proteomics of Ewing sarcoma tumors from patients following pembrolizumab therapy reveals upregulation of immunosuppressive proteins

Next, we sought to investigate changes in highly multiplexed protein expression on specific cell subsets following immunotherapy with pembrolizumab. Specifically, we investigated the expression of 68 proteins contained in the combined Bruke/Nanostring GeoMx immuno-oncology panels (**Table S1**) on a slide consecutive to slide utilized for the multiplexed immunofluorescence (Vectra) analysis. The GeoMx spatial proteomics platform permits isolation of proteins from full regions of interest, specific morphological regions, or individual cell populations. We examined protein expression in full regions of interest, stromal regions, and isolated CD8+ T cells and CD68+ macrophages (**Figure 3A**). Multiple ROIs were selected per tumor sample on the TMA to maximize material captured and included in the analysis. From the 68 proteins across these regions and cell types, we performed principal component analysis to evaluate whether there were consistent changes in protein levels between pre- and post-treatment with pembrolizumab (**Figure 3B**). These revealed that there was not strong separation between pre- and post-treatment across any of the regions analyzed, although there were some distinct regions enriched for pre-versus post-treatment samples from the stromal region and the full regions of interest.

**Figure 3.**
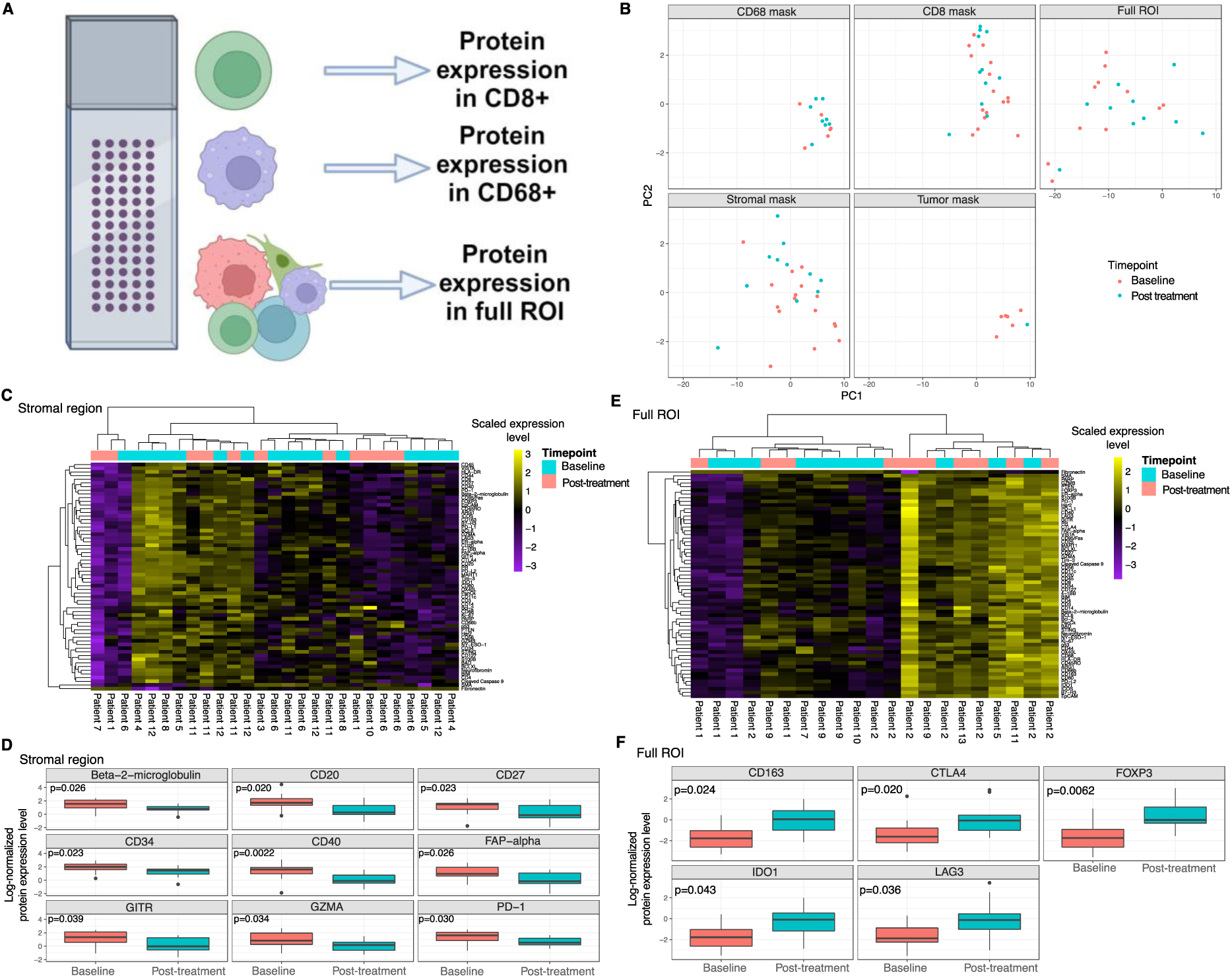
GeoMx spatial proteomic analysis reveals changes in the stromal regions and full regions of interest from baseline to post-treatment with pembrolizumab. **A)** Schematic depicting GeoMx analysis of specific cell types and regions. **B)** Principal component analysis of all the regions and masks collected by GeoMx. The stromal region and full regions of interest appear to have differences in the distribution and baseline and post-treatment samples. **C)** Heatmap showing protein expression profiles across the stromal mask from all patients. **D)** Significantly differentially expressed protein between pre- and post-treatment from the stroma region. Proteins associated with immune function (such as CD20, CD40, GITR, GZMA and PD-1) were lower in the stromal region following treatment. **E)** Heatmap showing protein expression profiles across full regions of interest from all patients. **F)** Significantly differentially expressed protein in full regions of interest show increase in immunoregulatory related proteins between baseline and post-treatment.

We next evaluated whether there were significantly differentially expressed proteins between baseline and post-treatment in the stromal region (**Figure 3C**). There was no clear separation between baseline and post-treatment samples by hierarchical clustering of gene expression profiles, but we did identify nine proteins that were differentially expressed in this region between baseline and post-treatment (**Figure 3D**). Many of these proteins are immune-related, such as CD20, GITR, GZMA, and PD-1, and had lower expression in the stromal region post-treatment, suggesting that pembrolizumab may alter immune composition or immune cell states in the stroma (versus exposure to pembrolizumab resulting in modulation of stromal-immune interactions). We also interrogated protein levels in full ROIs at baseline and post-treatment from the EwS TME (**Figure 3E**). There was some hierarchical clustering between pre- and post-treatment samples by protein expression in full ROIs (**Figure 3E**). This clustering pattern was at least partially driven by higher expression of immune regulatory proteins post-treatment, including CD163 (21), LAG3, FOXP3, CTLA4 and IDO1 (**Figure 3F**). We also interrogated whether there were correlations between protein abundance and percent change in tumor size across the stromal regions and the full regions of interest at either baseline or post-treatment. This analysis revealed that several features present in stromal regions of interest at baseline were significantly correlated with percent change in tumor size, although the limited number of evaluable patients preclude robust and generalizable conclusions (**Figure S4**). Overall, there were some changes in protein levels for immune-related features in the stromal region and full regions of interest following treatment with pembrolizumab, suggesting that there are some pharmacodynamic responses to treatment despite the absence of a clinical benefit.

### Joint integration approach maps cell types from single-cell RNAseq to CosMx

We next sought to evaluate single-cell expression profiles and physical locations of cells within tumors both before and after treatment with pembrolizumab. To achieve this, we utilized the Bruker/Nanostring CosMx Human Universal Cell Characterization RNA Panel (**Table S2**) to perform *in situ* single-cell spatial transcriptomics on our TMA (**Figure 4A-C**). After data processing and cell segmentation, we constructed gene expression profiles associated with individual cells (**Methods**). Next, we leveraged a single-cell RNAseq atlas of the Ewing sarcoma tumor microenvironment (including both treatment naïve and relapsed tumor specimens) (**Methods**) to create a reference atlas to serve as a robust map for interence of cell types in the CosMx spatial transcriptomics data. To achieve this, we first identified a shared latent space between these two heterogenous datasets (**Methods**). Briefly, we perform SCTransform-based integration and reciprocal principal component analysis to jointly embed the scRNAseq and CosMx data in the same space, facilitating label transfer from the fully annotated scRNAseq datasets to the CosMx dataset based on co-clustering (**Figure 4D; Methods**). This joint integration approach led to the robust annotation of single cells from the spatial transcriptomics dataset.

**Figure 4.**
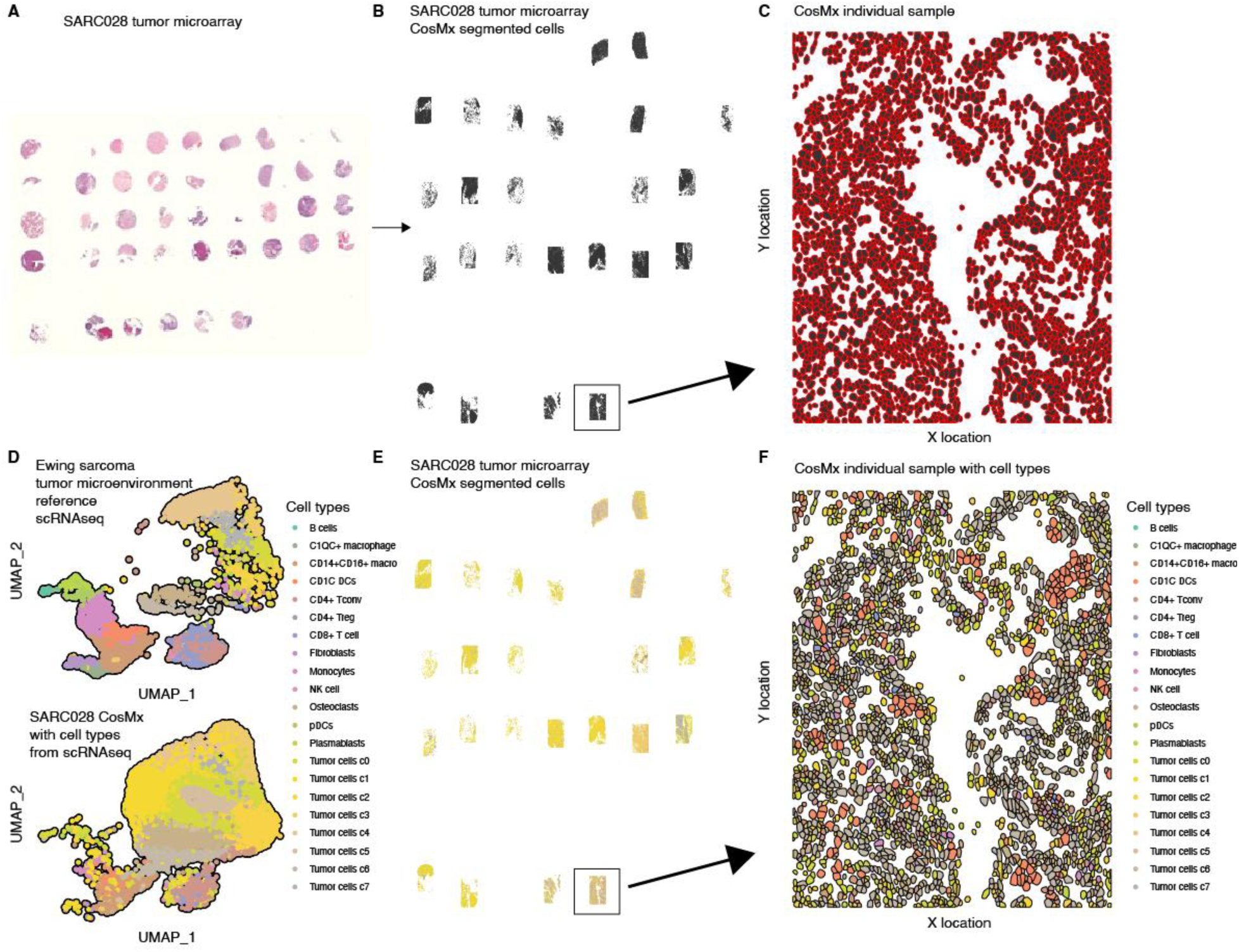
Joint integration of Ewing sarcoma single-cell RNAseq atlas and CosMx spatial gene expression profiling reveals spatial location of discrete cellular populations in the tumor microenvironment. **A)** Hematoxylin and eosin staining of a tumor microarray (TMA) generated from tissue biopsies obtained from SARC028 Ewing sarcoma patients at baseline and post-treatment with pembrolizumab. **B)** Same TMA as in **(A)**, but after imaging in the CosMx platform. **C)** Example of segmented cells obtained from one tumor from the TMA. **D)** Joint integration and clustering of a Ewing single-cell RNAseq atlas and the CosMx single-cell spatial transcriptomic profiles permitted robust inference of cell types based on CosMx RNA expression profiles. **E)** Same TMA image as in **(B)** but showing inferred cell types from the joint integration in **(D)**. **F)** Example of cell types and spatial location of segmented cells from the same individual tumor biopsy shown in **(C)**.

### Spatially resolved cellular neighborhoods are differentially associated with response to immunotherapy in Ewing sarcoma

After inferring cell types within our spatial transcriptomic dataset, we next evaluated whether there were recurrent spatial organizational patterns of cells in the EwS TME. To achieve this, we developed and implemented an unsupervised approach to identify spatially resolved cellular neighborhoods (CN; **Figure 5A**). Briefly, we identified the proportion of each cell type present within a given radius of a central cell (**Methods**). We repeated the process for all cells in the dataset and aggregated the vectors for each individual cell into a matrix. We then grouped these individual cell neighborhood vectors into clusters by performing K-means clustering. We note two important points about this approach: i) it identifies recurrent patterns present across patients; and ii) it is not reliant on any underlying distributional assumptions, which allows it to uncover abstract and irregularly shaped neighborhoods.

**Figure 5.**
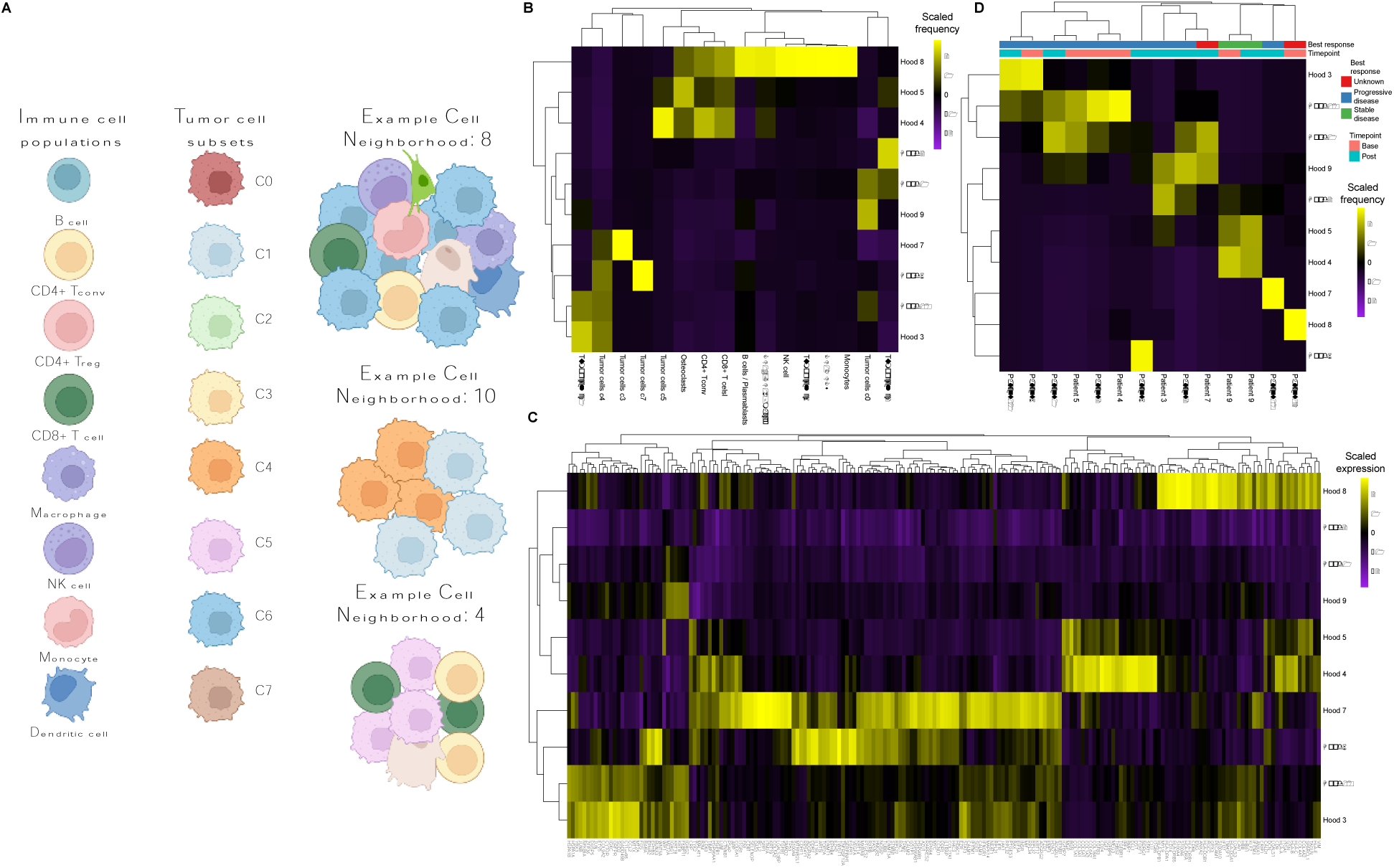
Identification of spatially resolved cellular neighborhoods in the Ewing tumor microenvironment and association with progressive disease. **A)** Schematic depicting examples of spatially resolved cellular neighborhoods identified by proximity based hierarchical clustering. **B)** Heatmap showing the enrichment of cell types in each of the 10 identified spatially resolved cellular neighborhoods. Both tumor subpopulations and immune subpopulations are differentially enriched across neighborhoods. **C)** Heatmap showing differentially expressed RNA across the cellular neighborhoods. **D)** Association between individual patient tumor biopsies and cellular neighborhoods. Cellular neighborhood 10 is consistently enriched at baseline and post-treatment in patients with progressive disease.

When we performed this analysis, we identified a total of 10 spatially resolved CNs (**Figure 5B**). These CNs were enriched for different cell types, with CNs 1, 2, 3, 6, 7, 9 and 10 predominantly consisting of tumor cell subsets. Specifically, CNs 3, 6, 7, and 10 consisted of mixtures of tumor subtypes while CNs 1, 2, and 9 consisted mostly of individual tumor cell subtypes. Conversely, CNs 4, 5 and 8 consisted of tumor cell types and non-tumor cell types including osteoclasts and immune cell subsets. CNs 4 and 5 were hierarchically related and were enriched for osteoclasts, CD8+ T cells and CD4+ T conventional cells. Neighborhood 8 was enriched for a diverse array of immune cells including the CD8+ and CD4+ T conventional cells present in neighborhoods 4 and 5 but also including other B lineage cells, myeloid cells, and NK cells. We also identified differentially expressed genes across each of these spatially resolved CNs. To achieve this, we leveraged the CN identities and performed a Wilcoxon rank sum test to compare neighborhoods in a one-versus-all based approach (**Methods**). This analysis revealed that these neighborhoods are representative of distinct biological states, reflected by the differentially expressed genes associated with each (**Figure 5C**). In summary, our approach for unsupervised identification of spatial patterns of cells in the EwS TME reveal specific neighborhoods of tumor cells as well as some neighborhoods that were enriched for either T cell or more diverse immune infiltrates.

We next questioned whether any CNs were consistently present across multiple patients. To achieve this, we performed hierarchical clustering of the proportion of cells in each patient that belonged to the CNs defined in **Figure 5B**. When we visualized the resulting heatmap, we also included the timepoint and response status for each individual patient, although we note that only one patient had stable disease and the remaining patients had progressive disease (**Figure 5D**). This analysis showed that CNs 4 and 5 were present at both baseline and post-treatment in Patient 9, the only patients that had stable disease. CNs 4 and 5 were associated with T cell infiltration and tumor cell subpopulation c5 (**Figure 5B**). Conversely, we found that CN10 was present in 6 patients that had progressive disease, suggesting that this neighborhood may be involved in resistance to anti-PD1 therapy. These results were further corroborated by principal component analysis (**Figure S5A-B**), which demonstrated that CN10 was a strong driver of separation along PC1. CN10 also constituted more than 50% of the cells in the tumor in patients that had high enrichment for CN10 (**Figure S5A**). CN10 was also consistently observed before and after treatment. CN10 mostly consisted of tumor cell subpopulations c1 and c4, suggesting that these tumor cell subtypes may have features the limit response to immunotherapy. We note that the small sample size precludes appropriately powered statistical analyses of these comparisons. Taken together, our cellular neighborhood approach revealed recurrent patterns of spatially resolved cellular neighborhoods, and a specific neighborhood (CN10) that was present across multiple patients with progressive disease.

### Dissection of tumor-specific neighborhoods reveals determinants of primary resistance to immunotherapy in Ewing sarcoma

After identifying that CN10 was associated with primary resistance to immunotherapy, we sought to better understand the features of this neighborhood that contribute to this resistance. To begin to understand the features associated with resistance to immunotherapy in EwS, we visualized the spatially resolved cellular neighborhoods in Patient 9 (patient with stable disease; **Figure 6A**) and Patient 2 (representative patient with progressive disease and high proportion of CN10; **Figure 6B**). This revealed, as anticipated, a high proportion of CN4 and 5 in Patient 9 (**Figure 6A**) and a high proportion of CN9 and CN10 in Patient 2 (**Figure 6B**). We also visualized the spatially resolved cellular neighborhoods across all patients and found that CN10 was significantly enriched across multiple patients (**Figure S6**). There were significant levels of immune infiltrate present in Patient 9, and little immune infiltrate in Patient 2 based on these cellular neighborhoods. We reasoned that the tumor cell subpopulations may be differentially contributing the immunogenicity of the TME, so we sought to determine whether there were differentially expressed genes across these tumor cell subpopulations (**Figure S7**). Tumor cell subpopulations c1 and c4 (enriched in CN10) were closely related by hierarchical clustering (**Figure S7**).

**Figure 6.**
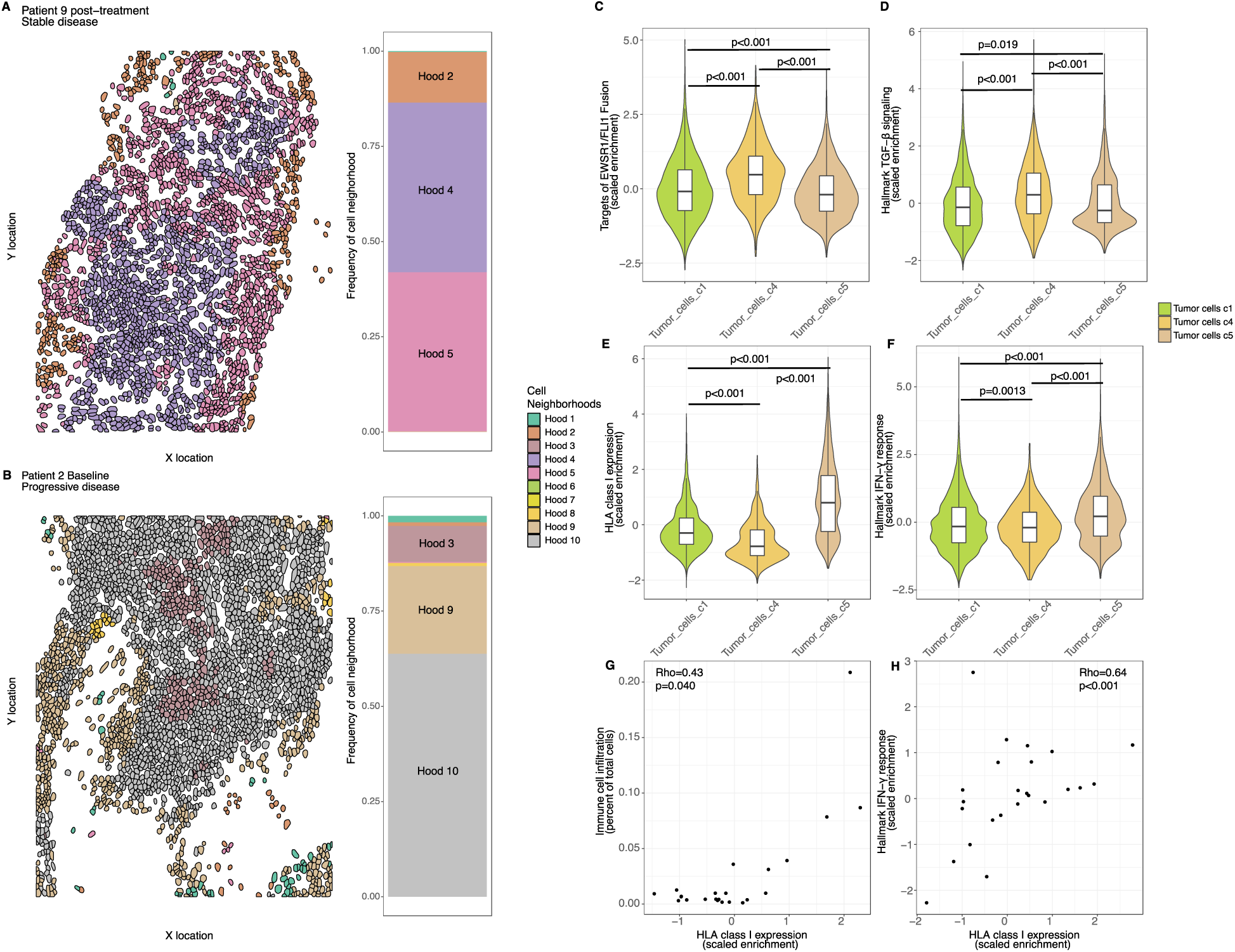
Low expression of HLA class I by specific tumor populations is associated with low type I interferon signaling, low immune infiltration, and progressive disease despite immunotherapy. **A)** Spatially resolved cellular neighborhoods 2, 4 and 5 are enriched in the patient that had stable disease following treatment with pembrolizumab. **B-E)** Gene set enrichment analysis of significantly different gene sets between tumor subpopulations c1 or c4 (enriched in cellular neighborhood 10 and associated with bad prognosis) versus tumor subpopulation c5 (enriched in cellular neighborhood 4 and associated with good prognosis). Tumor subpopulations c1 and c4 show higher EWS/FLI1 target expression **(B)** and TGF-β signaling **(C)** versus tumor subpopulation c5. Tumor subpopulation c5 shows higher expression of HLA class I **(D)** and IFN-γ signaling **(E)** versus tumor subpopulations c1 and c4. **G)** HLA class I expression and immune cell infiltration are correlated. **H)** HLA class I expression and IFN-γ signaling are strongly correlated.

From here, we sought to determine whether any gene sets were differentially expressed between the tumor subpopulations present in CN10 versus CNs 4 and 5. We first interrogated targets of EWS::FLI1, and found that tumor subpopulations c4 and to a lesser extent c1 (both enriched in CN10) had higher enriched of EWS::FLI1 targets versus tumor subpopulation c5 (enriched in CNs 4 and 5; **Figure 6C**). We also found that tumor subpopulation c4 and to a lesser extent c1 had significantly higher levels of TGF-β signaling versus tumor subpopulation c5 (**Figure 6D**), demonstrating that this immunosuppressive cytokine may also play a role in promoting the poor immunogenicity associated with CN10.

Given that antitumor immunity by CD8+ T cells is predicated on the expression of HLA class I molecules on the surface of tumor cells, we next interrogated the expression of HLA-A, HLA-B, and HLA-C across the tumor subpopulations present in CNs 4, 5, and 10. Surprisingly, this analysis revealed that tumor subpopulations c1 and c4 had dramatically lower expression of HLA class I molecules compared with tumor subpopulation c5 (which was enriched in the patient with stable disease following pembrolizumab). Based on the low levels of HLA class I expression, we next sought to further assess if this was potentially due to either the absence of interferon (IFN) responsiveness or other genomic defects. Correspondingly, we found that IFN-γ response signatures were higher in tumor subpopulation c5 versus c1 and c4. These findings are consistent with a model in which HLA class I expression is low due to impaired interferon response rather than genomic lesions that downregulate HLA expression. However, HLA class I expression has can be upregulated in EwS cells *in vitro* (22). To this end, we interrogate HLA class I expression on 3 EwS cell lines *in vitro* follow treatment with IFN-γ or control (**Figure S8**). Consistent with previous work, we find that HLA class I expression can be modified and upregulated in EwS. Overall, we identified several features of tumor subpopulations associated with CN10 that likely together contribute to reduce immune infiltration and inability to respond to immunotherapy.

## Discussion

To date, our understanding of the EwS TME, and factors in this microenvironment that predict immune infiltration of these tumors, has been limited. Here, we leveraged a unique opportunity to examine immune cell density, spatial proteomics, and single-cell spatial transcriptomics in Ewing tumor biopsy samples both before and after (8 weeks +/− 1 week) pembrolizumab therapy. We identify consistent CNs present in Ewing tumors associated with poor prognosis, as well as several potentially modifiable immunosuppressive features of these tumors that may contribute to the overall poor response to single agent checkpoint inhibition in this sarcoma subtype.

In patients enrolled on SARC028 with objective responses (those with dedifferentiated liposarcoma and undifferentiated pleomorphic sarcoma), there was an increase in immune cell populations following pembrolizumab therapy (12). Our finding confirms that in the setting of EwS in which there was poor response to single agent pembrolizumab, there is no effect on overall immune cell density between pre- and post-treatment biopsy samples. Tumor associated macrophages have been previously identified as a poor prognostic marker in Ewing tumors (5), and have increasingly been recognized to lead to CD8+ T cell inactivation. Interestingly, our analysis of differential protein expression following pembrolizumab therapy identified upregulation of CD163, a marker of an immunosuppressive macrophage population (23–26). We additionally identified an increase in expression of FOXP3, a marker of regulatory T cells, which, when present, are also known to blunt the activity of effector T cells (27). We hypothesize that the increased recruitment of these immunosuppressive cell populations to the EwS TME following pembrolizumab therapy may contribute to lack of clinical response. Future investigation of combinatorial therapies aimed at abrogating the immunosuppressive impact of macrophages and regulatory T cells in the EwS TME is warranted.

Mechanisms driving the net immunosuppression noted in relapsed EwS are still poorly understood. Recent scRNAseq studies of relapsed Ewing sarcoma, including our own work, have highlighted multiple mechanisms of immunosuppression in the EwS TME (6, 28). In our current study of paired patient samples from SARC028, we identify the triad of increased TGF-β signaling, reduced IFN response, and low expression of HLA type I as features of Ewing tumor cell populations that predominant in biopsies from patients with disease progression while receiving pembrolizumab. Here, we will consider these findings in the context of current data in the field. Previous work has demonstrated the important role of TGF-β signaling in EwS biology (29, 30). Inhibition of TGF-β has been clinically tested in patients with Ewing sarcoma including the use of TGF-β imprinted NK cellular therapy (NCT05634369) and as a component of VIGIL therapy which blocks TGF-β activation by reducing furin expression (NCT01061840). Our recent *in vivo* studies in humanized mice models of EwS have also demonstrated increased immune cell infiltration in EwS tumors following the combination of radiotherapy and inhibition of TGF-β signaling via a ligand trap (31). TGFβ inhibition is worthy of continued preclinical investigation in EwS.

EwS tumor cells have significant heterogeneity in response to cytokines in the TME such as interferon and TNF-α (22, 32). When considering interventions to ‘boost’ or ‘prime’ interferon responses in EwS, an area of active investigation in our laboratory, it is important to consider mechanisms to both induce interferon production in the EwS TME and to understand which Ewing tumors may be less responsive to interferon at baseline. Interferon response is critical given the ability of interferon stimulation to upregulate the expression key factors regulating immunotherapy response, such as tumor cell HLA class I expression (33). HLA class I expression in Ewing sarcoma is thought to be modifiable (22) in cells that retain robust responsiveness to inflammatory signaling. As previously suggested (28), a path forward for immunotherapy in the treatment of Ewing sarcoma is a focus on interventions that do not rely upon HLA class I expression for efficacy, such as targeted CAR T cell therapy (34). As immunosuppression in EwS is multifactorial, an alternative or complimentary approach is to utilize treatment combinations to block immunosuppression while promoting inflammation. Radiation therapy, a treatment modality commonly used to treat relapsed EwS, has the potential to both increase HLA class I expression and IFN response (35–37) and interventions to block immunosuppression specifically during radiation therapy in Ewing sarcoma warrant further investigation.

Lastly, this study highlights the importance of obtaining on-therapy biopsy samples as a component of clinical trials when feasible, especially for patients with rare tumor types such as Ewing sarcoma. Given the limited number of patients with Ewing sarcoma enrolled on SARC028, our current study could not determine differences in findings based on age, sex, or status of biologic factors such as STAG2 or TP53 mutation. It is increasingly clear that understanding tumor response to therapy, and in particular tumor characteristics that promote therapy resistance, is critical to designing new therapeutic approaches. Tumor heterogeneity is increasingly recognized as a critical pathway for tumor resistance to therapy (38). Our use of single-cell spatial transcriptomic data to identify cellular neighborhoods, with unique gene signatures and biologic states, highlights the heterogeneity of tumor cell populations and corresponding immune cell infiltration in EwS. Increased collection of Ewing biopsy samples in the setting of therapeutic trials or disease relapse is critical to advancing our understanding of the biology of this rare, heterogeneous, and aggressive cancer.

## Acknowledgements

The authors thank all members of the Cillo Lab and Bailey Lab for discussions and constructive feedback on the manuscript. We thank patients and their families for their willingness to consent to the acquisition of biopsy samples for research. We thank SARC for sharing biological specimens from the SARC028 trial. This research was supported in part by the University of Pittsburgh Center for Research Computing through the resources provided. Specifically, this work used the HTC cluster, which is supported by NIH award number S10OD028483. This work utilized the UPMC Hillman Cancer Center Cytometry Facility, a shared resource at the University of Pittsburgh supported by the CCSG P30 CA047904. This work was also supported by UPMC Hillman Cancer Center startup funds to the Cillo Lab and the Mario Lemieux Institute for Pediatric Research at UPMC Children’s (Bailey).

