## Supplemental Figures for "SARC028 samples reveal an interplay between TGF-beta, interferon signaling and low HLA class I expression as contributors to Ewing sarcoma checkpoint blockade resistance"

**Daley et al. Supplemental figures**

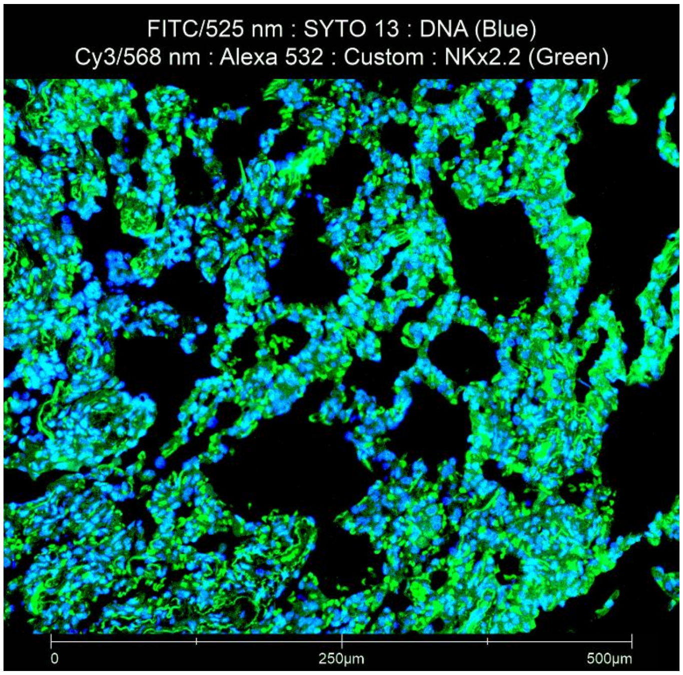


**Figure S1. NKX2.2 staining in Ewing tumor TMA samples.** Ewing tumor sample content was determined by NKX2.2 staining (green). DNA staining (blue) is used as a nuclei reference. Validation of NKX2.2 staining was conducted on formalin fixed, paraffin embedded (FFPE) human Ewing tumor specimens. NKX2.2 is used as the Ewing sarcoma tumor cell marker in GeoMx assays.

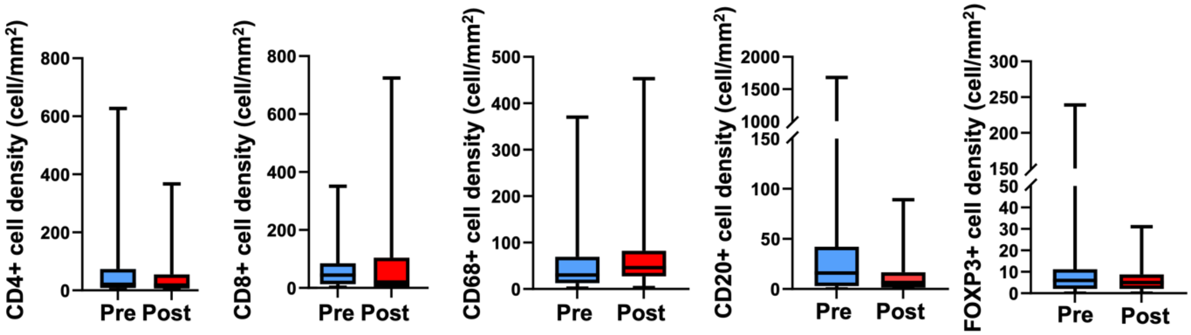


**Figure S2. Quantification of immune cell subsets utilizing multiplex immunohistochemistry.** Immune cell populations were not significant different across samples between pre- and post-pembrolizumab treatment.

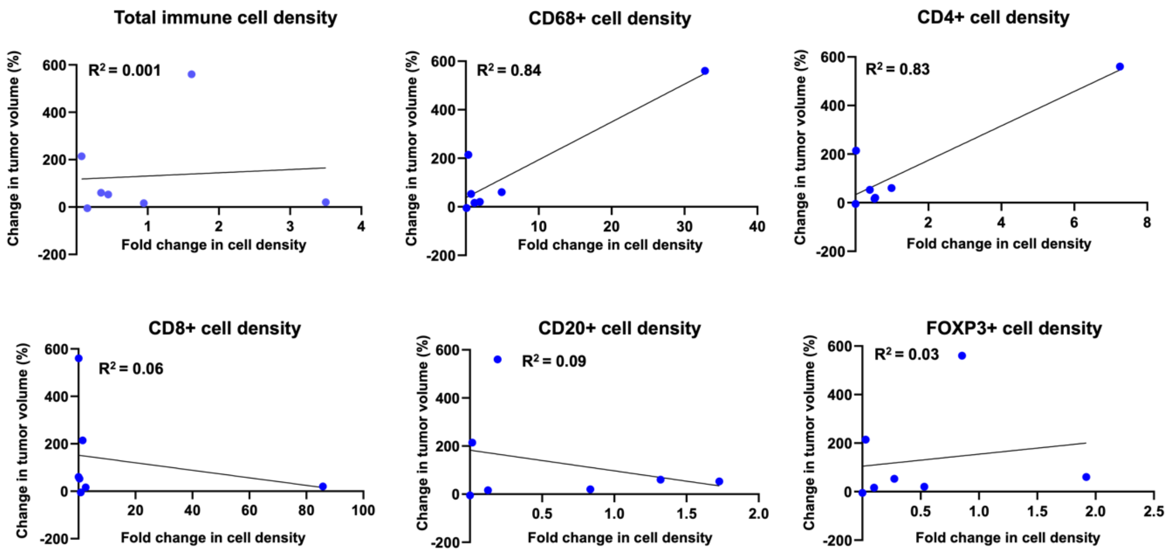


**Figure S3. Correlation between tumor response and fold change in immune cell infiltration.** Higher frequencies of CD68+ and CD4+ cell densities were associated with progressive disease (increased change in tumor volume between baseline and post-treatment).

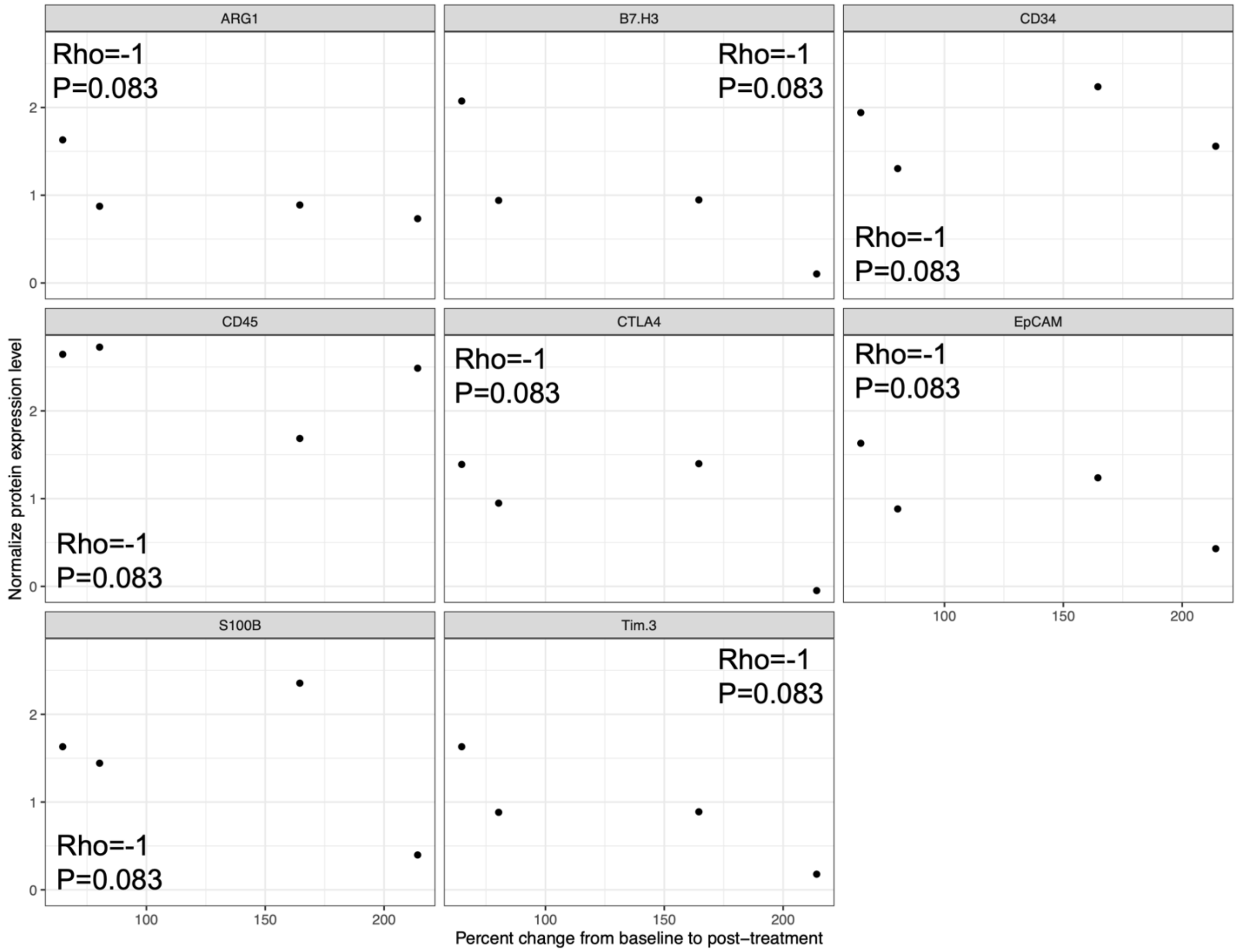


**Figure S4. Correlations between protein levels in stroma at baseline and changes in tumor size from pre- to post-treatment.** Levels of several proteins were negatively correlated with changes in tumor size between pre- and post-treatment, suggesting that these features are associated with better response.

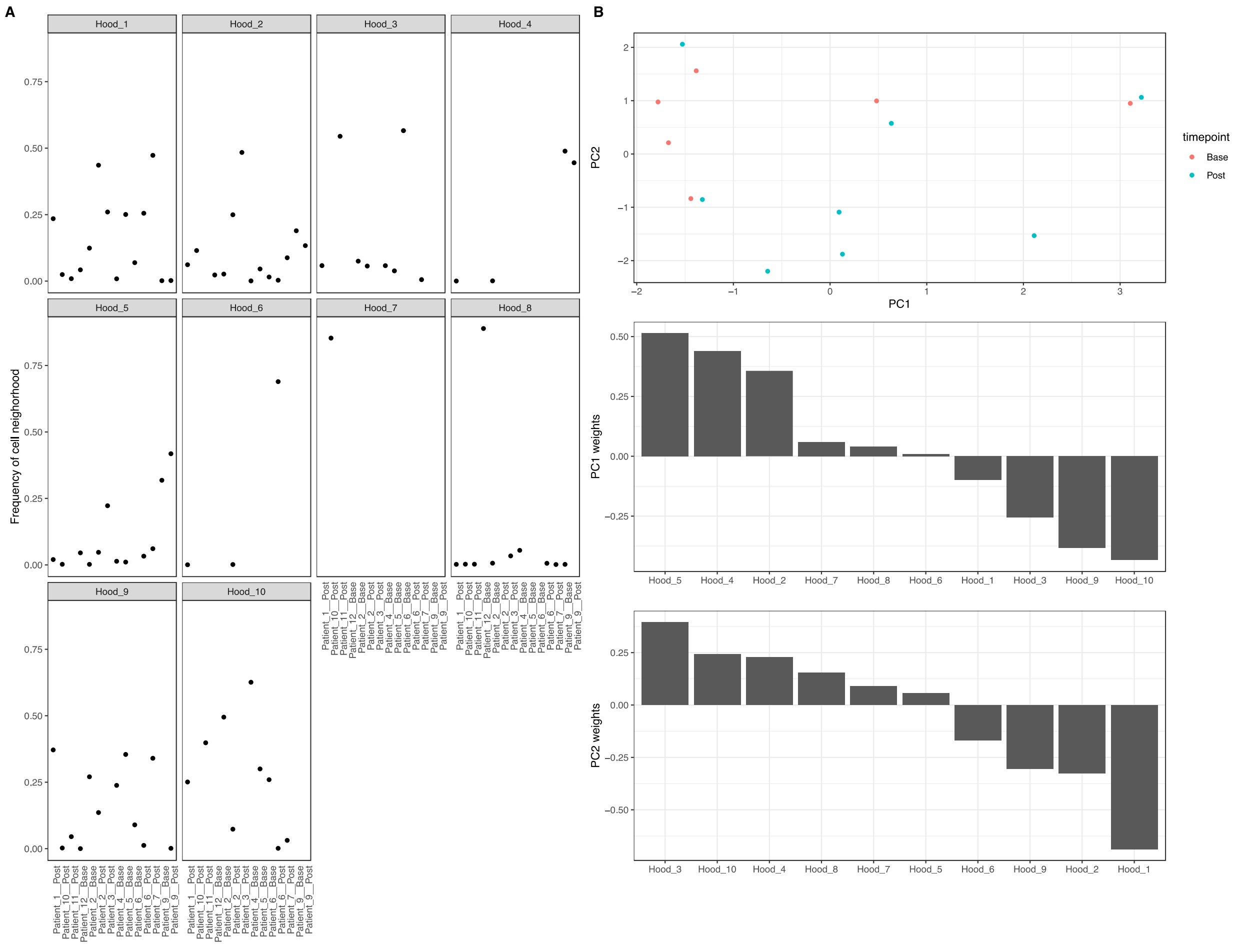


**Figure S5. Identification of groups of patients based on the frequency of spatially resolved cellular neighborhoods. A)** Quantification of the proportion of cells in each spatially resolved cell neighborhood across all specimens. **B)** Principal component analysis based on the frequency of cell neighborhoods across all patient specimens (top panel). Spatially resolved cellular neighborhoods that drive PC1 embeddings (middle panel) and PC2 embeddings (bottom panel). Notably, we observe a collection of patient samples with strong enrichment of spatially resolved cellular neighborhood 10 between -1 and -2 on the PC1 axis.

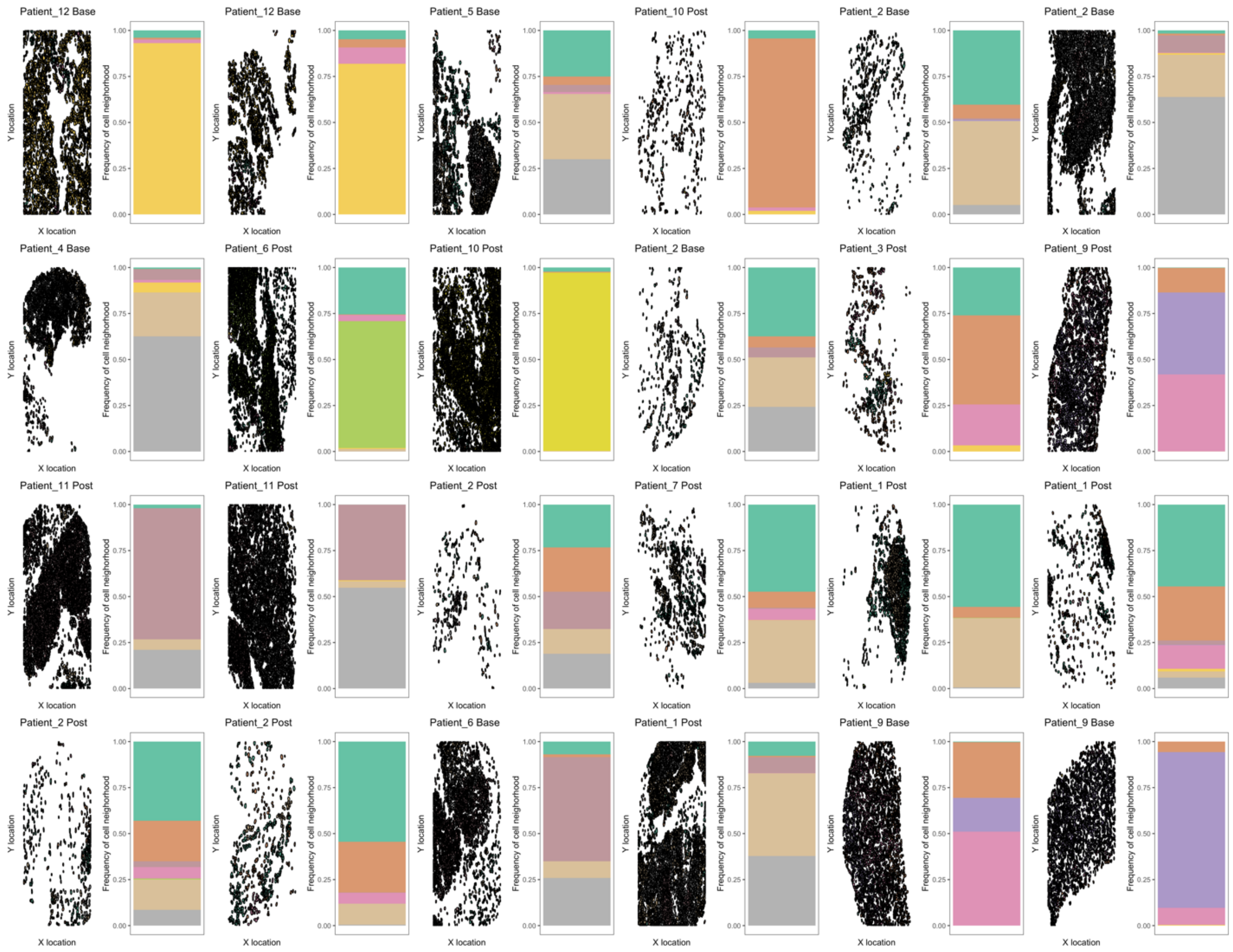


**Figure S6. Spatial distribution of cells from each patient tumor biopsy and enumeration of the proportion of cells with in each spatially resolved cell neighborhoods in each biopsy.**

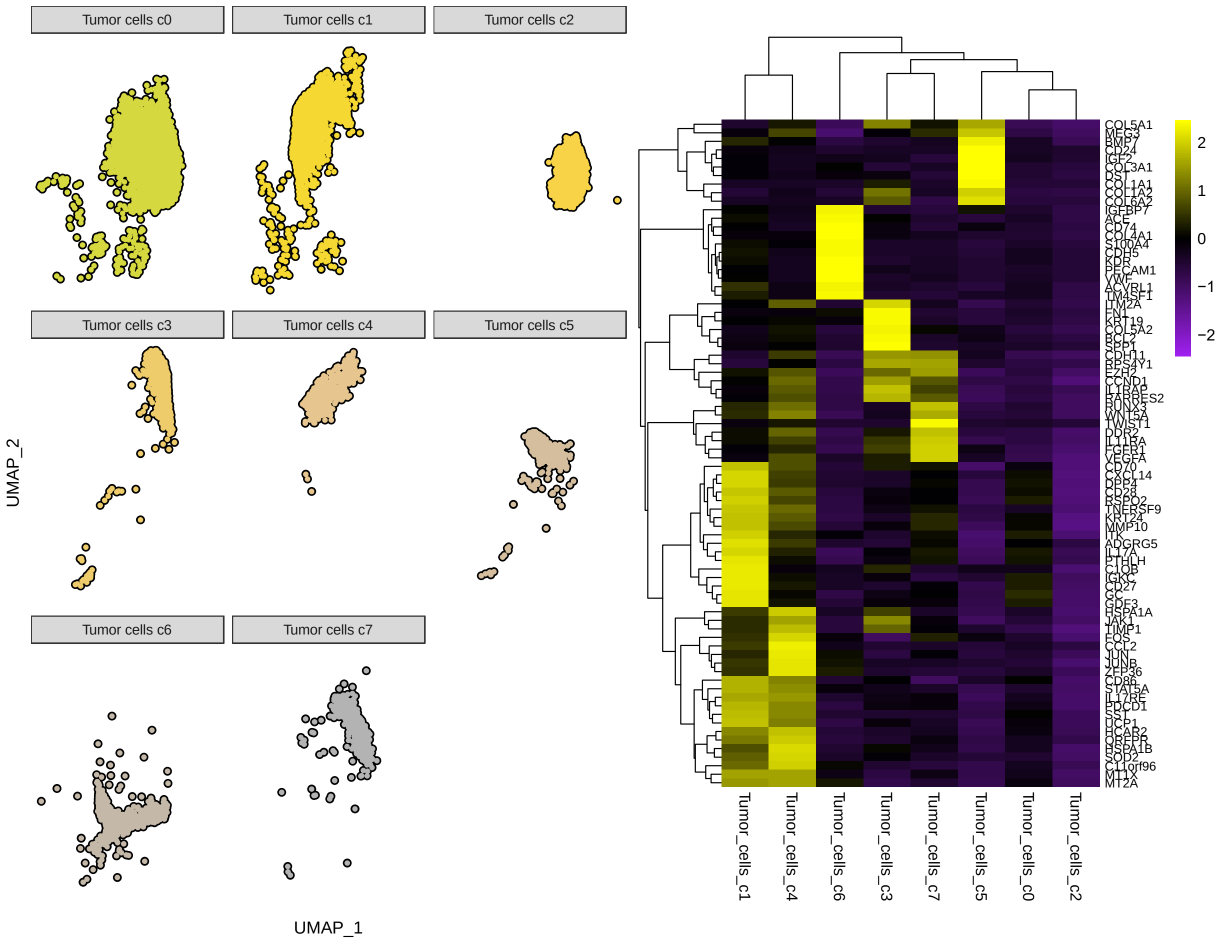


**Figure S7. Differentially expressed genes in Ewing sarcoma tumor cells associated with each cellular neighborhood. A)** UMAP embedding of all tumor subpopulations from the UMAP shown in **Figure 4**. **B)** Top 10 differentially expressed genes in each tumor subpopulation from the CosMx dataset.

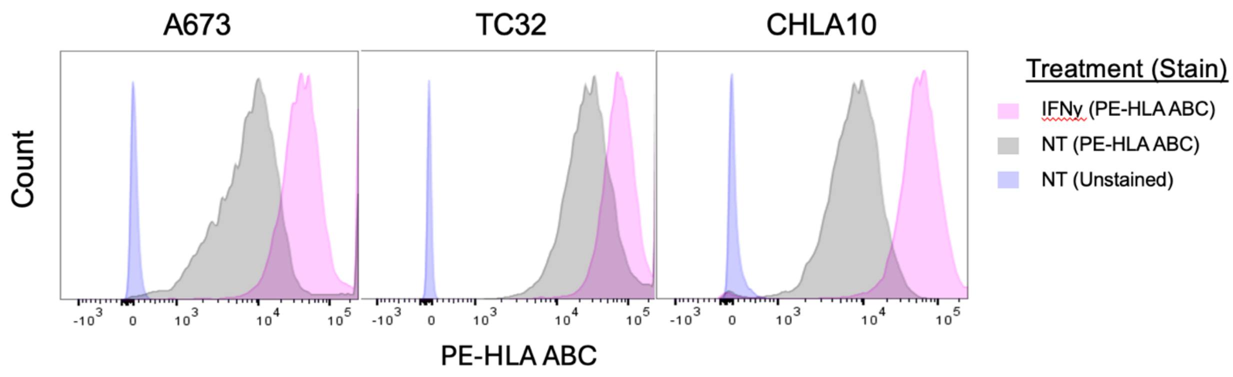


**Figure S8. HLA class I expression is modifiable and upregulated in Ewing sarcoma cells following interferon-gamma (IFNγ) treatment**. Ewing sarcoma cell lines (A673, TC32, CHLA10), were treated with 200 U IFNγ (or untreated control) for 16 hours. Surface expression of HLA-ABC determined by cell staining (PE-HLA ABC) or unstained controls and flow cytometry analysis (LSR Fortessa, 561nm laser). Data were analyzed and graphs generated utilizing FlowJo software v10.9.0.
